# Crystal structure of the 2019-nCoV spike receptor-binding domain bound with the ACE2 receptor

**DOI:** 10.1101/2020.02.19.956235

**Authors:** Jun Lan, Jiwan Ge, Jinfang Yu, Sisi Shan, Huan Zhou, Shilong Fan, Qi Zhang, Xuanling Shi, Qisheng Wang, Linqi Zhang, Xinquan Wang

## Abstract

A novel and highly pathogenic coronavirus (2019-nCoV) has caused an outbreak in Wuhan city, Hubei province of China since December 2019, and soon spread nationwide and spilled over to other countries around the world. To better understand the initial step of infection at atomic-level, we determined the crystal structure of the 2019-nCoV spike receptor-binding domain (RBD) bound with the cell receptor ACE2 at 2.45 Å resolution. The overall ACE2-binding mode of the 2019-nCoV RBD is nearly identical to that of the SARS-CoV RBD, which also utilizes ACE2 as the cell receptor. Structural analysis identified residues in 2019-nCoV RBD critical for ACE2 binding, and majority of which are either highly conserved or shared similar side chain properties with those in the SARS-CoV RBD. Such similarity in structure and sequence strongly argue for a convergent evolution between 2019-nCoV and SARS-CoV RBD for improved binding to ACE2 despite of being segregated in different genetic lineages in the betacoronavirus genus. The epitopes of two SARS-CoV antibodies targeting the RBD are also analyzed with the 2019-nCoV RBD, providing insights into future identification of cross-reactive antibodies.

---

The emergence of a novel and highly pathogenic coronavirus (2019-nCoV) in Wuhan city, Hubei province of China and its rapid international spread has posed a serious global public health emergency^1–3^. Similar to those infected by pathogenic severe acute respiratory syndrome coronavirus (SARS-CoV) in 2003 and Middle East respiratory syndrome coronavirus (MERS-CoV) in 2012, patients infected by 2019-nCoV manifested a range of symptoms including dry cough, fever, headache, dyspnea and pneumonia with estimated mortality rate of 2.5%^4–6^. Since the initial outbreak in December of 2019, 2019-nCoV has spread throughout China and to more than twenty other countries worldwide. As of February 17, 2020, 70641 cases in China have been confirmed with the infection while 7264 cases are suspected, and 1772 cases have died. Currently, the epicenter Wuhan and the neighboring cities have been under lockdown to minimize continued spread, and the WHO has announced a Public Health Emergency of International Concern (PHEIC) duo to the rapid and global dissemination of 2019-nCoV.

Phylogenetic analysis on the coronavirus genomes has revealed that 2019-nCoV is a new member of the betacoronavirus genus, which includes SARS-CoV, MERS-CoV, bat SARS-related coronaviruses (SARSr-CoV), as well as others identified in humans and diverse animal species^1–3,7^. Bat coronavirus RaTG13 appears to be the closest relative of the 2019-nCoV sharing over 93.1% homology in the spike (S) gene. SARS-CoV and other SARSr-CoVs however are rather distinct with less than 80% homology^1^.

Coronaviruses utilize the homotrimeric spike glycoprotein (S1 subunit and S2 subunit in each spike monomer) on the envelope to bind their cellular receptors. Such binding triggers a cascade events leading to the fusion between cell and viral membranes for cell entry. Our cryo-EM studies have shown that the binding of SARS-CoV spike to the cell receptor ACE2 induces the dissociation of the S1 with ACE2, prompting the S2 to transition from a metastable prefusion to a more stable postfusion state that is essential for membrane fusion^8,9^. Therefore, binding to ACE2 receptor is a critical initial step for SARS-CoV to entry into the target cells. Recent studies also pointed to the important role of ACE2 in mediating entry of 2019-nCoV^1,10^. HeLa cells expressing ACE2 is susceptible to 2019-nCoV infection while those without failed to do so^1^. In vitro SPR experiments also showed that the binding affinity of ACE2 to the spike glycoprotein and to the receptor-binding domain (RBD) are equivalent, with the former of 14.7 nM and the latter of 15.2 nM^11,12^. These results indicate that the RBD is the key functional component within the S1 subunit responsible for binding to ACE2.

The cryo-EM structure of the 2019-nCoV spike trimer at 3.5 Å resolution has just been reported^12^. The coordinates are not yet available for detailed characterization. However, inspection of the structure features presented in the uploaded manuscript on bioRxiv indicated incomplete resolution of RBD in the model, particularly for the receptor-binding motif (RBM) that interacts directly with ACE2. Computer modeling of interaction between 2019-CoV RBD and ACE2 has identified some residues potentially involved in the actual interaction but the actual interaction remained elusive^13^. Furthermore, despite of impressive cross-reactive neutralizing activity from serum/plasma of SARS-CoV recovered patients^14^, no SARS-CoV monoclonal antibodies targeted to RBD so far isolated are able to bind and neutralize 2019-nCoV^11,12^. These findings highlight some intrinsic sequence and structure differences between the SARS-CoV and 2019-nCoV RBDs.

To elucidate the 2019-nCoV RBD and ACE2 interaction at a higher resolution, we chose to determine the complex structure of 2019-nCoV RBD bound with ACE2 by X-ray crystallography. The atomic-level structural information would greatly improve our understanding of interaction between 2019-nCoV and susceptible cells, providing precise target for neutralizing antibodies, and assisting structure-based vaccine design urgently needed in our ongoing combat against 2019-nCoV. Specifically, we expressed the 2019-nCoV RBD (residues Arg319-Phe541) (Fig. 1a and 1b) and the N-terminal peptidase domain of ACE2 (residues Ser19-Asp615) in Hi5 insect cells and purified them by Ni-NTA affinity and gel-filtration (Fig. S1). The complex structure was determined by molecular replacement using the SARS-CoV RBD and ACE2 structures as search models, and refined at 2.45 Å resolution to final *R*_work_ and *R*_free_ factors of 21.9% and 27.8%, respectively (Fig. S2 and Table S1). The final model contains residues Cys336 to Glu516 of 2019-nCoV RBD and residues Ser19 to Asp615 of ACE2 N-terminal peptidase domain, as well as 63 solvent molecules.

**Fig. 1.**
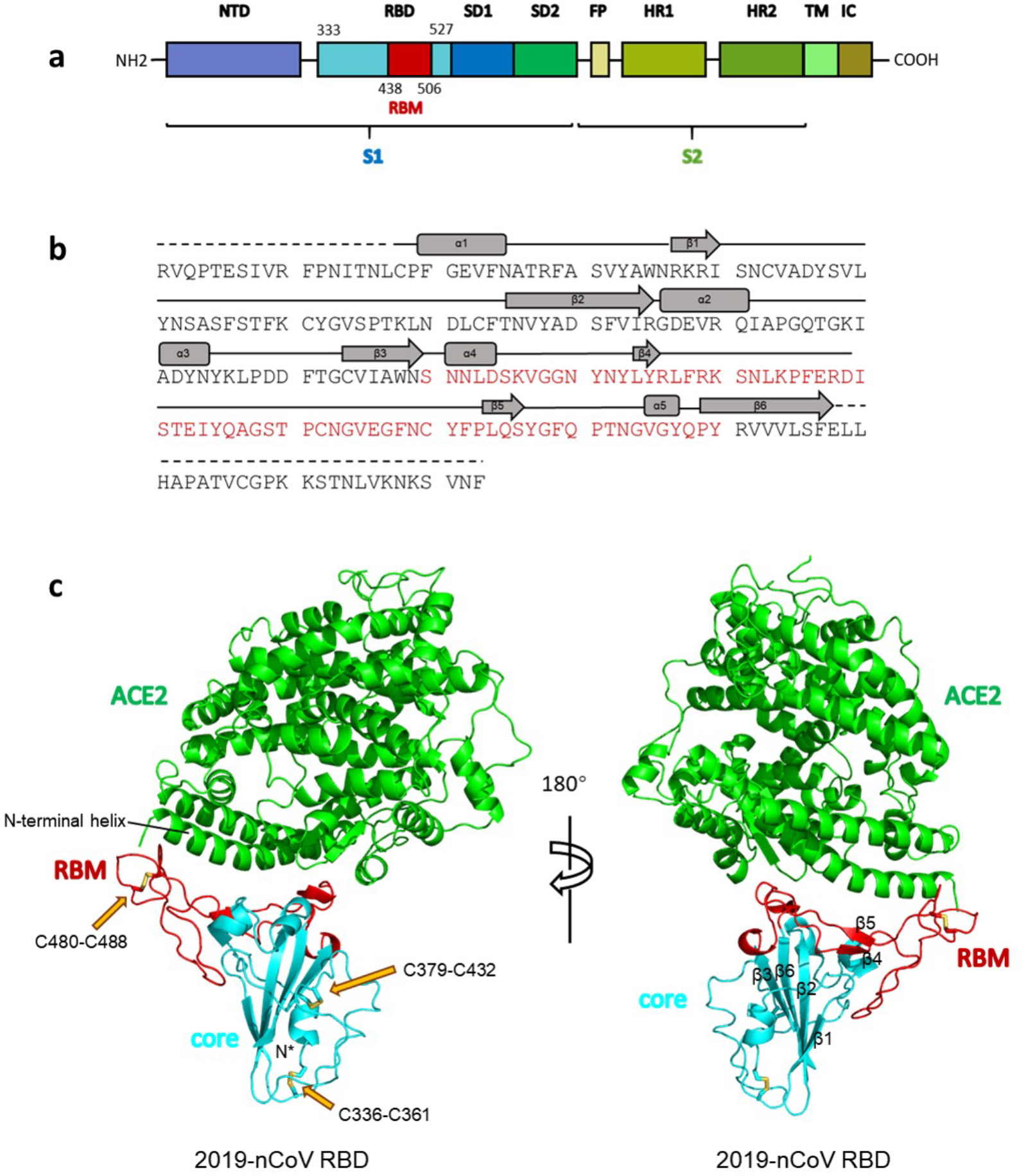
Overall structure of 2019-nCoV RBD bound with ACE2. (**a**) Overall topology of 2019-nCoV spike monomer. NTD, N-terminal domain. RBD, receptor-binding domain. RBM, receptor-binding motif. SD1, subdomain 1. SD2, subdomain 2. FP, fusion peptide. HR1, heptad repeat 1. HR2, heptad repeat 2. TM, transmembrane region. IC, intracellular domain. (**b**) Sequence and secondary structures of 2019-nCoV RBD. The RBM is colored red. (**c**) Overall structure of 2019-nCoV RBD bound with ACE2. ACE2 is colored green. 2019-nCoV RBD core is colored cyan and RBM is colored red. Disulfide bonds in the 2019-nCoV RBD are shown as stick and indicated by yellow arrows. The N-terminal helix of ACE2 responsible for binding is labeled.

The 2019-nCoV RBD has a twisted four-stranded antiparallel β sheet (β1, β2, β3 and β6) with short connecting helices and loops forming as the core (Fig. 1b and 1c). Between the β3 and β6 strands in the core, there is an extended insertion containing short β4 and β5 strands, α4 and α5 helices and loops (Fig. 1b and 1c). This extended insertion is the receptor-binding motif (RBM) containing most of the contacting residues of 2019-nCoV for ACE2 binding. A total of nine cysteine residues are found in the RBD, six of which forming three pairs of disulfide bonds are resolved in the final model. Among these three pairs, two are in the core (Cys336-Cys361 and Cys379-Cys432) to help stabilize the β sheet structure (Fig. 1c) while the remaining one (Cys480-Cys488) connects loops in the distal end of the RBM (Fig. 1c). The N-terminal peptidase domain of ACE2 has two lobes, forming the peptide substrate binding site between them. The extended RBM in the 2019-nCoV RBD contacts the bottom side of the ACE2 small lobe, with a concave outer surface in the RBM accommodating the N-terminal helix of the ACE2 (Fig. 1c). The overall structure of the 2019-nCoV RBD is similar to that of the SARS-CoV RBD (Fig. 2a), with an r.m.s.d. of 1.2 Å for 174 aligned Cα atoms. Even in the RBM that has more sequence variations, the overall structure is also highly similar (r.m.s.d. of 1.3 Å) with only one obvious conformational change in the distal end (Fig. 2a). The overall binding model of the 2019-nCoV RBD to the ACE2 is also nearly identical to that observed in previously determined SARS-CoV RBD/ACE2 complex structure^15^ (Fig. 2b).

**Fig. 2.**
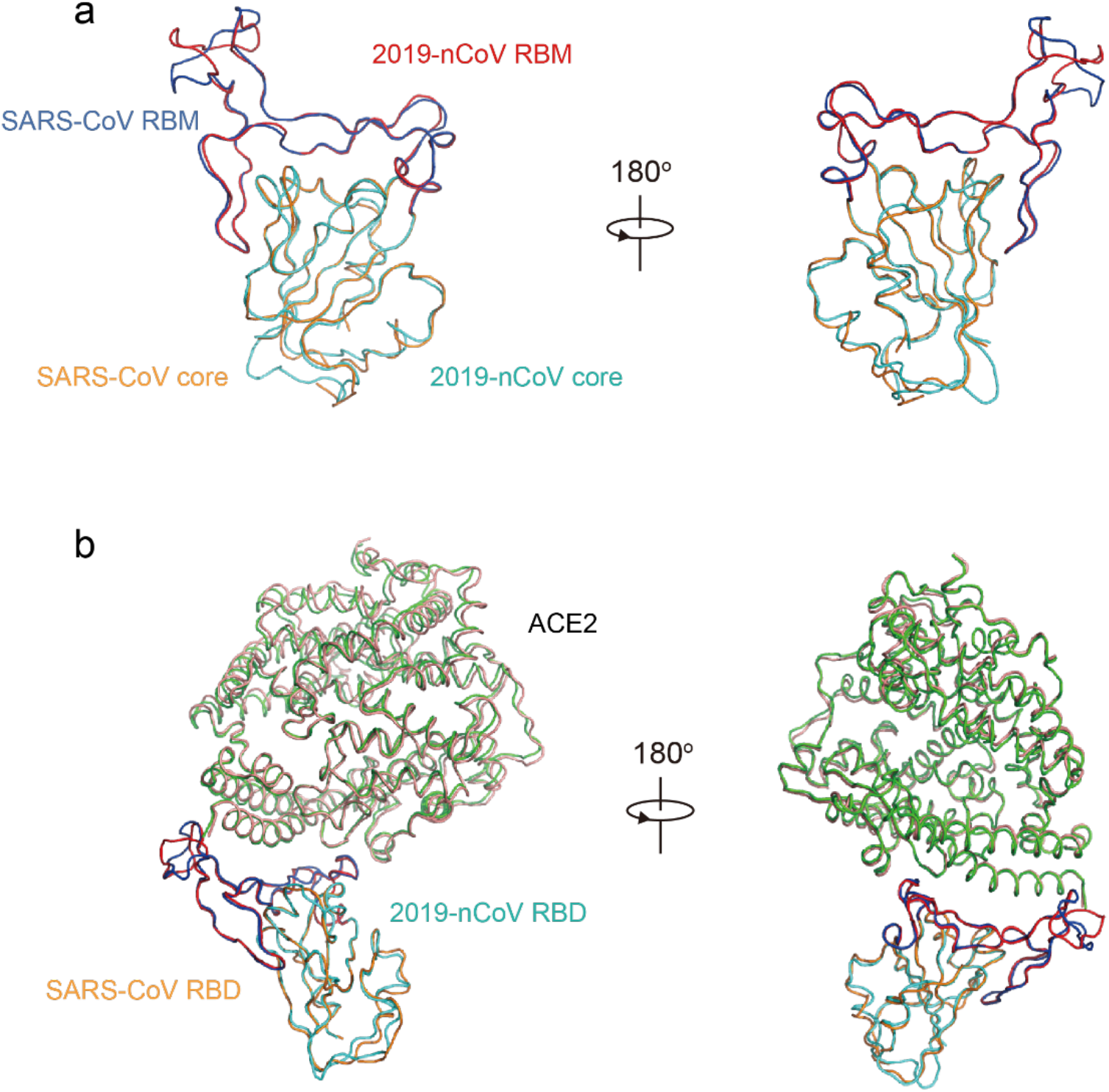
Structural comparisons of 2019-nCoV and SARS-CoV RBDs and their binding modes to ACE2 receptor. **(a)** Alignment of the 2019-nCoV RBD (core in cyan and RBM in red) and SARS-CoV RBD (core in orange and RBM in blue) structures. **(b)** Structural alignment of 2019-nCoV RBD/ACE2 and SARS-CoV RBD/ACE2 complexes. 2019-nCoV RBD is colored cyan and red, its interacting ACE2 is colored green. SARS-CoV RBD is colored orange and blue, its interacting ACE2 is colored salmon. The PDB code for SARS-CoV RBD/ACE2 complex: 2AJF.

The cradling of the ACE2 N-terminal helix by the RBM outer surface results in a large buried surface of ~1700 Å^2^ between the 2019-nCoV RBD and ACE2 receptor (Fig. 1c). With a distance cutoff of 4 Å, a total of 18 residues of the RBD contact 20 residues of the ACE2 (Fig. 3a and Table S2). Analysis of interface between SARS-CoV RBD and ACE2 revealed a total of 16 residues of the SARS-CoV RBD contact 20 residues of the ACE2 (Fig. 3a and Table S2). Among the 20 residues interacting with the two different RBDs, 17 are shared and most of which are located at the N-terminal helix (Fig. 2a). One prominent and common feature presented at both interfaces is the networks of hydrophilic interactions. There are 17 hydrogen bonds and 1 salt bridge at the 2019-nCoV RBD/ACE2 interface, and 12 hydrogen bonds and 2 salt bridges at the SARS-CoV RBD/ACE2 interface (Table 1). Another shared feature is the involvement of multiple tyrosine residues in forming hydrogen-bonding interactions with the polar hydroxyl group. These include Tyr449, Tyr489, Tyr495 and Tyr505 from the 2019-nCoV RBD and Tyr436, Tyr475 and Tyr491 from the SARS-CoV RBD (Table 1). To further identify and compare the ACE2-interacting residues, we used structure-guided sequence alignment and mapped them onto their respective RBD sequences (Fig. 3b). Among 14 shared amino acid positions utilized by both RBMs for ACE2 interaction, eight have the identical residues between the 2019-nCoV and SARS-CoV RBDs including Tyr449/Tyr436, Tyr453/Tyr440, Asn487/Asn473, Tyr489/Tyr475, Gly496/Gly482, Thr500/Thr486, Gly502/Gly488 and Tyr505/Tyr491 (Fig. 3b). Five positions have residues demonstrating similar biochemical properties despite of having different side chains including Leu455/Tyr442, Phe456/Leu443, 486Phe/Leu472, Gln493/Asn479 and Asn501/Thr487 (Fig. 3b). Four of the five SARS-CoV residues such as Tyr442, Leu472, Asn479 and Thr487 have previously been shown to be critical for ACE2 binding^13^. The remaining one is at the Gln498/484Tyr position, while the SARS-CoV RBD Tyr484 is not involved in hydrogen-bonding interaction, the Gln498 of the 2019-nCoV RBD forms hydrogen-bonding interaction with Gln42 of ACE2 (Table 1). Outside RBM, there is a unique ACE2-interacting residues Lys417 in the 2019-nCoV, forming a salt-bridge with ACE2 Asp30 (Fig. 3b). This position is replaced by a valine in the SARS-CoV RBD that fails to participate in ACE2 binding (Fig. 3b). Consistently, comparison of the surface electrostatic potential also identified a positive-charged patch on the 2019-nCoV RBD contributed by Lys417 that is absent on the SARS-CoV RBD (Fig. 3c). Taken together, these results show that the 2019-nCoV RBD/ACE2 and SARS-CoV RBD/ACE2 interfaces share substantial similarity in the buried surface area, the number of interacting residues, and hydrophilic interaction networks although some differences in surface electrostatic potential were identified (Fig. 3). Such similarity argues strongly for the convergent evolution of the 2019-nCoV and SARS-CoV RBD structures to improve binding affinity to the same ACE2 receptor despite of being in different genetic lineages in the betacoronavirus genus.

**Table 1.** The hydrogen bonds and salt bridges identified using PISA program at the 2019-nCoV RBD/ACE2 and SARS-CoV RBD/ACE2 interfaces.

|  | 2019-nCoV RBD | Distance(Å) | ACE2 | Distance(Å) | SARS-CoV RBD |
| --- | --- | --- | --- | --- | --- |
| Hydrogen bonds | N487(ND2) | 2.4 | Q24(OE1) | 2.9 | N473(ND2) |
|  | Q493(NE2) | 3.2 | E35(OE1) |  |  |
|  | **Y505(OH)** | 3.5 | E37(OE1) | 3.4 | **Y491(OH)** |
|  | **Y505(OH)** | 3.6 | E37(OE2) |  |  |
|  | **Y449(OH)** | 3.0 | D38(OD1) | 3.0 | **Y436(OH)** |
|  | **Y449(OH)** | 2.8 | D38(OD2) | 3.0 | **Y436(OH)** |
|  | T500(OG1) | 2.7 | Y41(OH) | 2.8 | T486(OG1) |
|  | N501(N) | 3.7 | Y41(OH) | 3.3 | T487(N) |
|  | Q498(OE1) | 3.1 | Q42(NE2) |  |  |
|  | G446(O) | 3.6 | Q42(NE2) |  |  |
|  | **Y449(OH)** | 3.2 | Q42(NE2) |  |  |
|  |  |  | Q42(OE1) | 2.7 | **Y436(OH)** |
|  | N487(OD1) | 2.5 | Y83(OH) | 2.8 | N473(ND2) |
|  | **Y489(OH)** | 3.4 | Y83(OH) | 3.3 | **Y475(OH)** |
|  |  |  | Q325(OE1) | 3.8 | R426(NH2) |
|  |  |  | N330(ND2) | 2.8 | T486(O) |
|  | **Y495(O)** | 3.5 | K353(NZ) |  |  |
|  | G496(O) | 3.1 | K353(NZ) |  |  |
|  | G502(N) | 2.7 | K353(O) | 2.5 | G488(N) |
|  | **Y505(OH)** | 3.8 | R393(NH2) |  |  |
| Salt bridges | K417(NZ) | 3.1 | D30(OD2) |  |  |
|  |  |  | E329(OE2) | 3.7 | R426(NH1) |
|  |  |  | E329(OE2) | 2.9 | R426(NH2) |

**Fig. 3.**
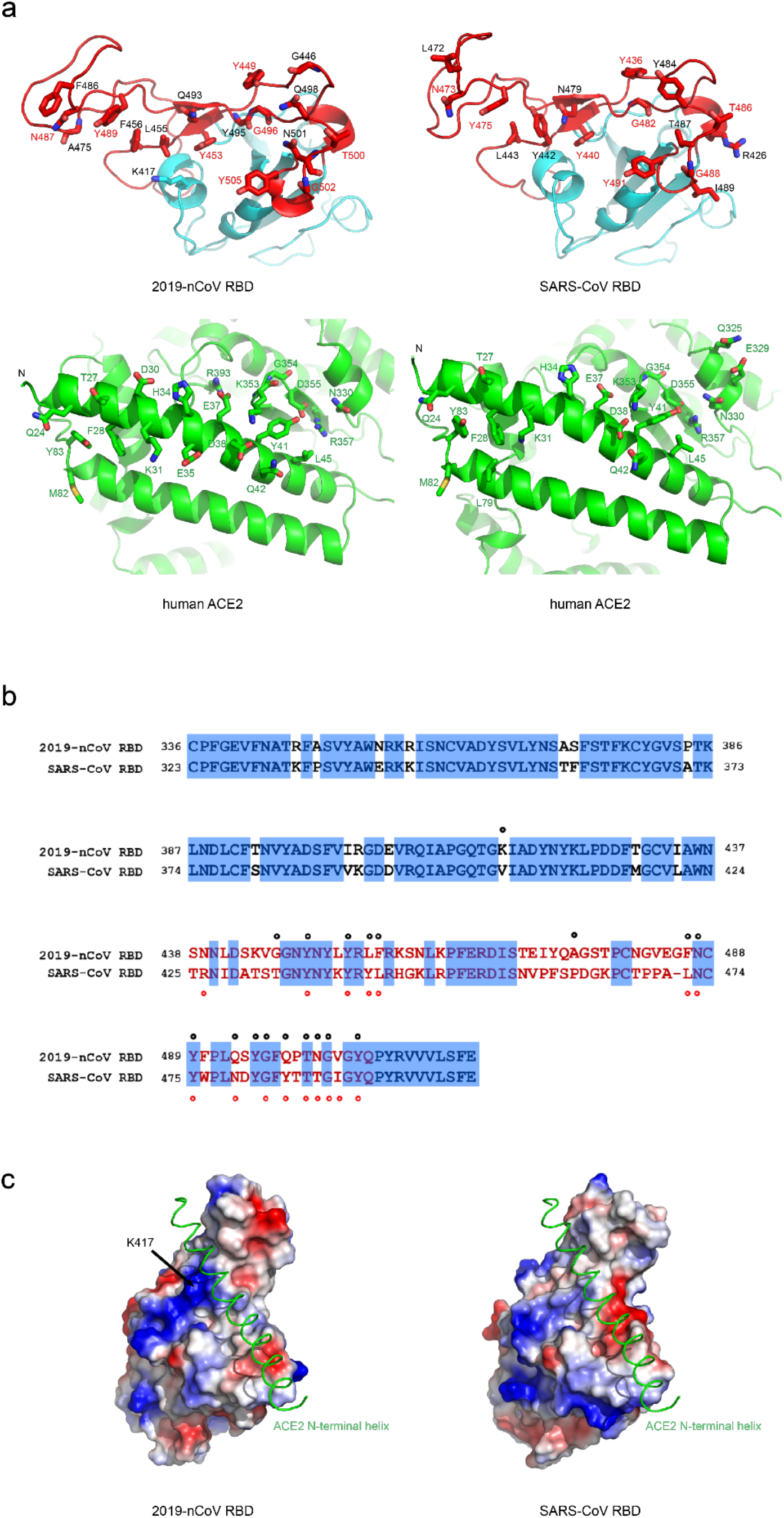
The 2019-nCoV RBD/ACE2 binding interface compared with that of SARS-CoV RBD/ACE2. **(a)** Contacting residues shown as stick at the 2019-nCoV RBD/ACE2 and SARS-CoV RBD/ACE2 interfaces. Positions in both RBDs involved in ACE2 binding are indicated by red labels. **(b)** Sequence alignment of 2019-nCoV RBD and SARS-CoV RBD. Contacting residues in the 2019-nCoV RBD are indicated by black dots; contacting residues in the SARS-CoV RBD are indicated by red dots. **(c)** Electrostatic potential map of 2019-nCoV RBD (left panel), and SARS-CoV RBD (right panel). The position of K417 in the 2019-nCoV RBD is indicated by black arrow. The N-terminal helix of ACE2 is shown as green ribbon. The PDB code for SARS-CoV RBD/ACE2 complex: 2AJF.

Consistent with structural similarity, the binding affinities between ACE2 and 2019-nCoV and SARS-CoV RBDs also fall into the same range (~10-60 nM) as previously reported^11,12^. However, this is somewhat different from a recent report where an ~10-20 fold increased binding between ACE2 and 2019-nCoV spike trimer was found (K_D_ of 14.7 nM) compared with that between ACE2 and SARS-CoV RBD-SD1 (K_D_ of 325 nM)^12^. This is perhaps due to the different proteins used in the assay or some other unknown reasons. Nevertheless, the binding affinity alone is unlikely to explain the unusual transmissibility of 2019-nCoV. Other factors such as the unique “RRAR” furin cleavage site at the S1/S2 boundary of the 2019-nCoV spike may play more important roles in facilitating the rapid human-to-human transmission.

Neutralizing antibodies represent a critical component of immune system in fighting against viral infection. It has been reported that the 2019-nCoV could be cross-neutralized by horse anti-SARS-CoV serum and convalescent serum from SARS-infected patient^1,14^, reinforcing structural similarity between 2019-nCoV and SARS-CoV RBDs. Such similarity also raised the hope of rapid application of previously characterized SARS-CoV monoclonal antibodies in the clinical setting. However, no antibody targeted to SARS-CoV (m396, S230, 80R and CR3014) has so far demonstrated any impressive cross-binding and neutralization activity against 2019-nCoV spike or RBD^11,12,16–19^. One exception is SARS-CoV antibody CR3022 that binds to the 2019-nCoV RBD with a K_D_ of 6.2 nM, although its neutralizing activity against 2019-nCoV has not been reported yet^11^. Currently, we are uncertain where exactly the epitope of CR3022 on SARS-CoV nor on 2019-nCoV RBDs. Among the three antibodies incapable of binding to the 2019-nCoV RBD, two (m396 and 80R) have the epitopes resolved by high resolution crystal structure determination of SARS-CoV RBD-Fab complexes. Through mapping these epitope residues onto the sequence of SARS-CoV RBD aligned with the sequence of 2019-nCoV RBD (Fig. 4), we found that antibody m396 has seven residue changes in the 2019-nCoV RBD among 21 epitope positions (Fig. 4). There are 15 reside changes in the 2019-nCoV RBD among 24 epitope positions by antibody 80R (Fig. 4). This may provide the structural basis for the lack of cross-reactivity by m396 and 80R. However, conserved residues between 2019-nCoV and SARS-nCoV RBD indeed exist, even in the more variable RBM (Fig. 4). The cross-neutralization of 2019-nCoV by horse anti-SARS-CoV serum and serum/plasm from recovered SARS patients indicates a great potential in identifying antibodies with cross-reactivity between these two coronaviruses^1,14^. Such antibody will present a great promise for developing therapeutic agents toward diverse coronavirus species including 2019-nCoV.

**Fig. 4.**
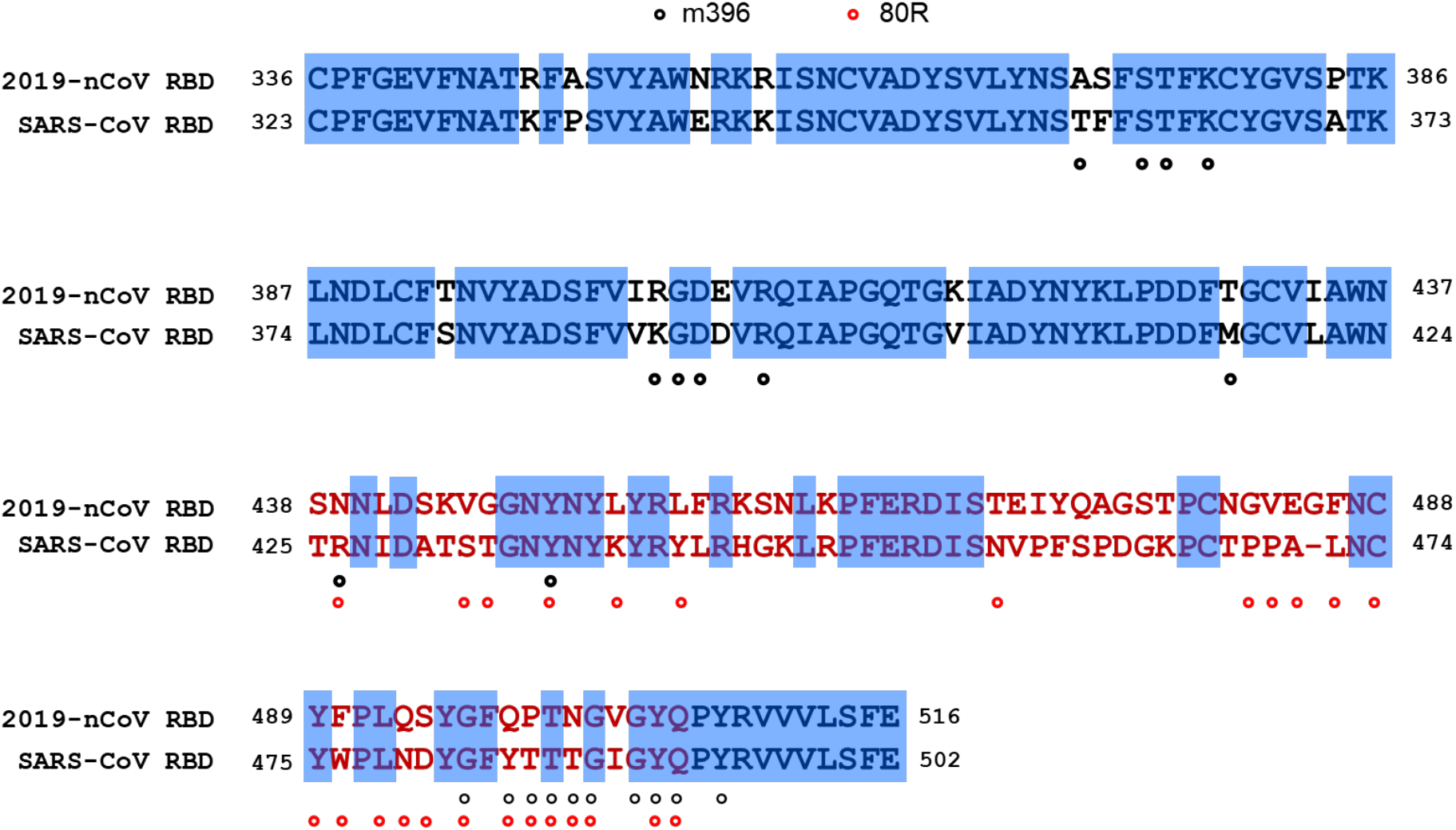
Mapping of SARS-CoV neutralizing antibody epitopes. The epitopes of SARS-CoV neutralizing antibodies m396 and 80R, which target the RBD, are labeled in the SARS-CoV sequence aligned with the sequence of 2019-nCoV RBD. Epitope residues of m396 are indicated by black dots; epitope residues of 80R are indicated by red dots.

## Materials and Methods

### Protein expression and purification

The 2019-nCoV receptor-binding domain (RBD) and the N-terminal peptidase domain of human ACE2 were expressed using the Bac-to-Bac baculovirus system (Invitrogen). The 2019-nCoV RBD (residues Arg319-Phe541) with an N-terminal gp67 signal peptide for secretion and a C-terminal 6×His tag for purification was inserted into pFastBac-Dual vector (Invitrogen). The construct was transformed into bacterial DH10Bac component cells, and the extracted bacmid was then transfected into Sf9 cells using Cellfectin II Reagent (Invitrogen). The low-titer viruses were harvested and then amplified to generate high-titer virus stock, which was used to infect Hi5 cells at a density of 2×10^6^ cells/ml. The supernatant of cell culture containing the secreted RBD was harvested 60 h after infection, concentrated and buffer-exchanged to HBS (10 mM HEPES, pH 7.2, 150 mM NaCl). RBD was captured by Ni-NTA resin (GE Healthcare) and eluted with 500 mM imidazole in HBS buffer. RBD was then purified by gel filtration chromatography using the Superdex 200 column (GE Healthcare) pre-equilibrated with HBS buffer. Fractions containing RBD were collected.

The N-terminal peptidase domain of human ACE2 (residues Ser19-Asp615) was expressed and purified by essentially the same protocol used for the 2019-nCoV RBD. To purify the 2019-nCoV RBD/ACE2 complex, ACE2 was incubated with RBD for 1 h on ice in HBS buffer, and the mixture was then subjected to gel filtration chromatography. Fractions containing the complex were pooled and concentrated to 13 mg/ml.

### Crystallization and data collection

Crystals were successfully grown at room temperature in sitting drops, over wells containing 100 mM MES, pH 6.5, 10% PEG5000mme, 12% 1-propanol. The drops were made by mixing 200 nL RBD/ACE2 in 20 mM Tris pH 7.5, 150 mM NaCl with 200 nL well solution. Crystals were harvested, soaked briefly in 100 mM MES, pH 6.5, 10% PEG5000mme, 12% 1-propanol, 20% glycerol, and flash-frozen in liquid nitrogen. Diffraction data were collected at the BL17U beam line of the Shanghai Synchrotron Research Facility (SSRF)^20^. Diffraction data were auto-processed with aquarium pipeline and the data processing statistics are listed in Table S1^21^.

### Structural determination and refinement

The structure was determined by the molecular replacement method with PHASER in CCP4 suite^22^. The search models are ACE2 extracellular domain and SARS-CoV RBD (PDB code 2AJF). Density map improvement by atoms update and refinement was performed with ARP/wARP^23^. Subsequent model building and refinement were performed using COOT and PHENIX, respectively^24,25^. The structural refinement statistics are listed in Table S1. All structural figures were generated with PyMol^26^.

## Acknowledgments

We thank the SSRF BL17U beam line for data collection and processing. We thank at the X-ray crystallography platform of the Tsinghua University Technology Center for Protein Research for providing the facility support. This work was supported by funds from Beijing Advanced Innovation Center for Structural Biology at Tsinghua University and the National Key Plan for Scientific Research and Development of China (grant number 2016YFD0500307). It is also supported by Tsinghua University Initiative Scientific Research Program (20201080053), the National Natural Science Foundation Award (81530065), Beijing Municipal Science and Technology Commission (171100000517-001 and −003), and Tencent Foundation, Shuidi Foundation, and TH Capital.

---

**Fig. S1.**
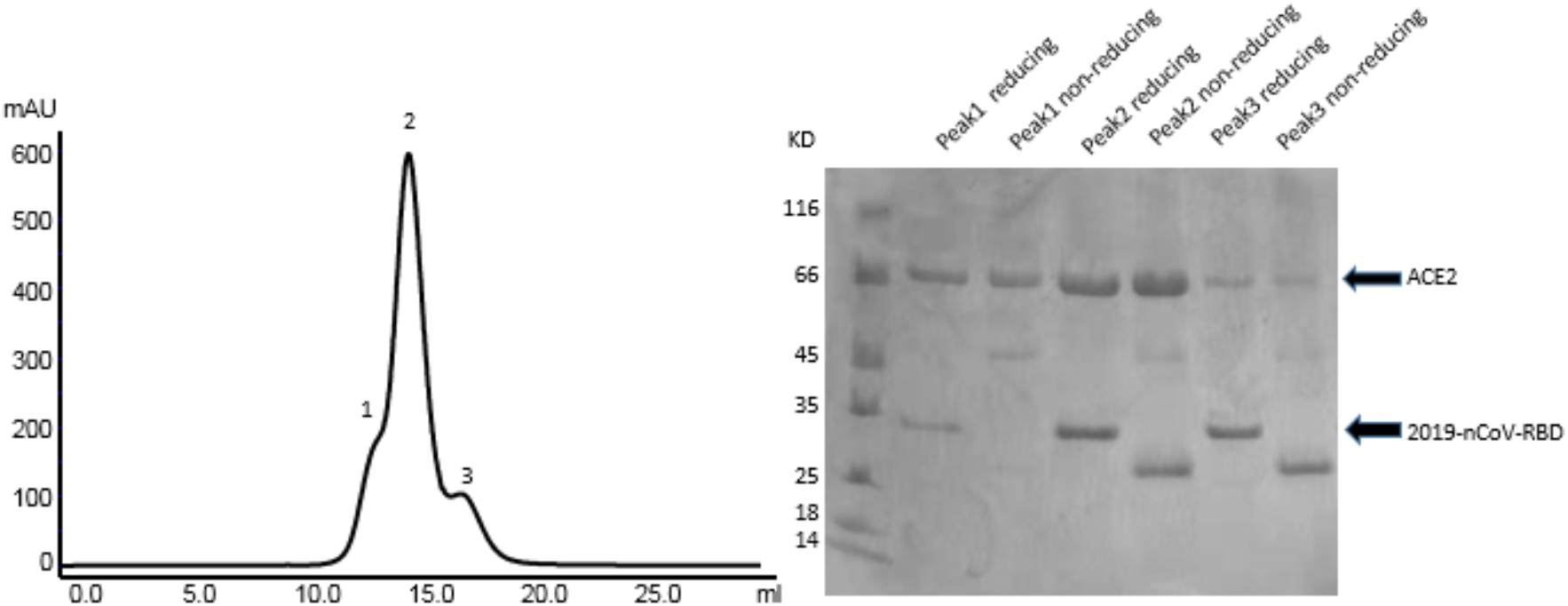
Purification of 2019-nCoV RBD/ACE2 complex

**Fig. S2.**
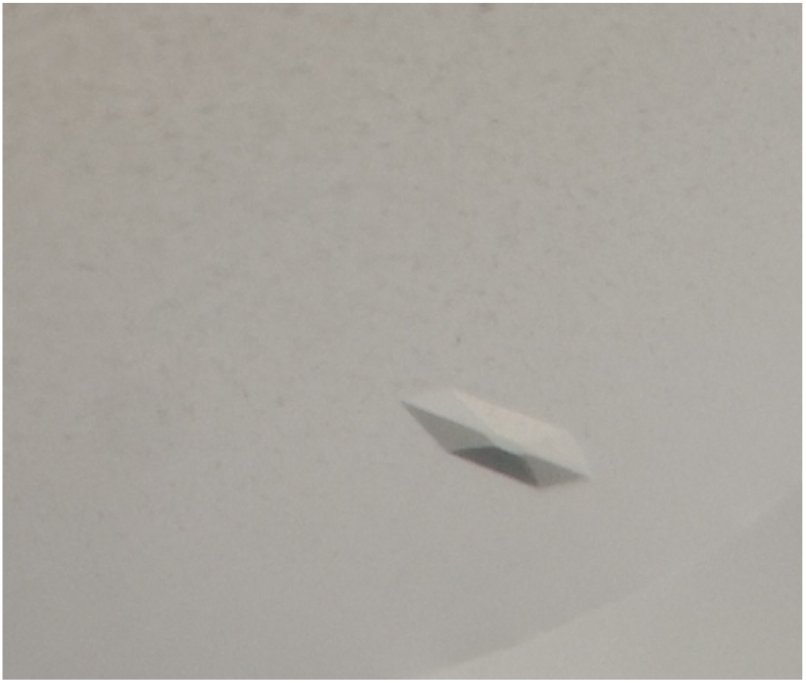
Crystal of 2019-nCoV RBD/ACE2 complex.

**Table S1.** Data collection and refinement statistics.

| **Data Collection** |  |
| --- | --- |
| Beamline | SSRF BL17U |
| Wavelength | 0.9796 Å |
| Space group | P4<sub>1</sub>2<sub>1</sub>2 |
| Cell dimensions |  |
| a, b, c (Å) | 104.67, 104.67, 228.72 |
| α, β, γ (°) | 90, 90, 90 |
| Resolution (Å) | 53.1- 2.45 (2.54 - 2.45) |
| <sup>a</sup>R<sub>merge</sub> | 0.118 (2.70) |
| <sup>b</sup>R<sub>pim</sub> | 0.033 (0.738) |
| <sup>c</sup>CC<sub>1/2</sub> of the highest resolution shell | 0.661 |
| I / σI | 24.2 (1.7) |
| Completeness (%) | 99.90 (99.98) |
| Redundancy | 26.1 (27.3) |
| **Refinement** |  |
| Resolution (Å) | 53.1- 2.45 |
| No. Reflections | 47545 |
| <sup>d</sup>R<sub>work</sub> / R<sub>free</sub> (%) | 21.1/23.7 |
| No. atoms |  |
| Protein | 6369 |
| Water | 16 |
| B-factors (Å<sup>2</sup>) |  |
| Protein | 68.2 |
| Water | 58.8 |
| r.m.s. deviations |  |
| Bond lengths (Å) | 0.014 |
| Bond angles (°) | 1.90 |
| Ramachandran plot (%) |  |
| Most favored | 95.01 |
| Allowed | 4.61 |
| Disallowed | 0.38 |

<sup>a</sup>R<sub>merge</sub> = $\sum_{hkl} \sum_j |I_j(hkl) - \langle I(hkl) \rangle| / \sum_{hkl} \sum_j I_j(hkl)$ , where *I* is the intensity of reflection.
<sup>b</sup>R<sub>pim</sub> = $\sum_{hkl} [I/(N-I)]^{1/2} \sum_j |I_j(hkl) - \langle I(hkl) \rangle| / \sum_{hkl} \sum_j I_j(hkl)$ , where *N* is the redundancy of the dataset.
<sup>c</sup>CC<sub>1/2</sub> is the correlation coefficient of the half datasets.
<sup>d</sup>R<sub>work</sub> = $\sum_{hkl} |F_{obs} - F_{calc}| / \sum_{hkl} |F_{obs}|$ , where *F<sub>obs</sub>* and *F<sub>calc</sub>* is the observed and the calculated structure factor, respectively. *R<sub>free</sub>* is the cross-validation *R* factor for the test set of reflections (5% of the total) omitted in model refinement.

**Table S2.** Contact residues at the 2019-nCoV RBD/ACE2 and SARS-CoV RBD-ACE2 interfaces

| 2019-nCoV RBD-ACE2 interface |  | SARS-CoV-ACE2 interface |  |
| --- | --- | --- | --- |
| ACE2 | 2019-nCoV RBD | ACE2 | SARS-CoV RBD |
| 24Q | 417K | 24Q | 426R |
| 27T | 446G | 27T | 436Y |
| 28F | 449Y | 28F | 440Y |
| 30D | 453Y | 31K | 442Y |
| 31K | 455L | 34H | 443L |
| 34H | 456F | 37E | 472L |
| 35E | 475A | 38D | 473N |
| 37E | 486F | 41Y | 475Y |
| 38D | 487N | 42Q | 479N |
| 41Y | 489Y | 45L | 482G |
| 42Q | 493Q | 79L | 484Y |
| 45L | 495Y | 82M | 486T |
| 82M | 496G | 83Y | 487T |
| 83Y | 498Q | 325Q | 488G |
| 330N | 500T | 329E | 489I |
| 353K | 501N | 330N | 491Y |
| 354G | 502G | 353K |  |
| 355D | 505Y | 354G |  |
| 357R |  | 355D |  |
| 393R |  | 357R |  |

